# Newly-identified lentil genotypes adapted to Mediterranean agro-ecosystems

**DOI:** 10.1101/2025.07.10.664142

**Authors:** Lorenzo Rocchetti, Alex Kumi Frimpong, Valerio Di Vittori, Chiara Santamarina, Alice Pieri, Andrea Tosoroni, Simone Papalini, Francesca Francioni, Evan Musari, Elisa Bellucci, Laura Nanni, Stefania Marzario, Giuseppina Logozzo, Tania Gioia, Kirstin Bett, Guido Arlotti, Marco Silvestri, Roberto Papa, Elena Bitocchi

## Abstract

Lentil cultivation and consumption promote human health and sustainable agriculture, making a significant contribution to the transition toward a plant-based diet. In Europe, lentil yields are still unstable, and the lack of breeding efforts limits the choice of farmers to few varieties. Here, we characterized 46 lentil genotypes, including local cultivars and landraces from diverse geographic origins, in Mediterranean agro-environments for flowering, architectural and production traits in seven field trials, over 3 years (2019-2021), in two localities (central and southern Italy) and during two sowing seasons (autumn and spring). We estimated the genetic merit of each genotype and identified outperforming genotypes for all traits. Indian ILL 11557AGL and Argentinian IL 4605AGL domesticated varieties resulted superior for earliness. Italian landraces and French cultivars achieved the highest values for first pod height, while landraces and breeding materials from Ethiopia, Syria and Iran were the best-yielding. Data from all seven trials were available for 16 genotypes, so we analyzed the genotype, environment and genotype × environment interaction (GEI) to identify specific genotypic adaptations. European cultivars performed well for architectural traits, whereas the best-yielding genotypes were Middle Eastern and Ethiopian landraces. Environmental effect on yield related to sowing season and locality was detected, with an overall higher yield in autumn compared to spring sowing trials and in central rather than southern Italy. By dissecting the GEI structure using additive main effect and multiplicative interaction (AMMI) analysis and Weighted Average of Absolute Scores (WAASB) index, we identified a group of Iranian landraces (PI 431633 AGL and PI 432033 LSP AGL) adapted to both autumn and spring sowing and one Ethiopian landrace (IG 1959 AGL) showing high yield stability across all environmental conditions. These findings provide a foundation to unlock the full potential of lentil cultivation in European and Mediterranean systems by identifying adapted, high-performing genotypes.

## 1. Introduction

Feeding the growing global population while reducing the environmental impact of the industrial agriculture and mitigating the effects of climate change, requires a major shift toward a plant-based diet (Poore and Nemecek, 2018; Shukla et al., 2019; Gerten et al., 2020). In this context, food legumes offer a valuable protein source as an alternative to meat (Bellucci et al., 2021), and an important source of minerals, complex carbohydrates, dietary fiber and vitamins, as well as byproducts that can be exploited as sources of bioactive compounds and ingredients in a circular economy (Nartea et al., 2023). Legumes not only produce nutritious seeds, they also establish symbiotic associations with nitrogen-fixing bacteria, helping to increase soil nitrogen levels when used in rotation with other crops and/or as a green manure crop, thus enhancing agro-ecosystem productivity (Dabin et al., 2016; Lindström and Mousavi, 2019; Gerten et al., 2020; Aguilar et al., 2022). This also reduces the environmental impact of industrial agriculture by reducing dependence on nitrogen fertilizers produced using non-renewable fossil resources (Paris et al. 2022). Accordingly, increasing the cultivation of food legumes in Europe would confer significant ecosystem benefits by regulating the nutrient cycle and reducing greenhouse gas emissions (Watson et al., 2017).

Lentil (*Lens culinaris* Medik.) is a traditional legume crop in Europe and in the Mediterranean Basin. It was domesticated in the Fertile Crescent, probably southern Turkey (Zohary, 1972; Ladizinsky, 1979; Alo et al., 2011), during the Neolithic period and subsequently spread into Europe, Africa, and south Asia (Sonnante et al., 2009). In 2022, global lentil production reached 6.6 Mt from 5.5 Mha of land. The leading producer is Canada (2.30 Mt), followed by India (1.26 Mt) and Australia (0.99 Mt), whereas Europe is only a minor producer (0.26 Mt) (FAOSTAT 20220, assessed on 12 October 2024).

In Italy, lentil production is based mainly on landraces grown on marginal fields with relatively low yield potential (Laghetti et al., 2008). Increases in lentil production and expansion to different environments are hindered by the limited number of available cultivars, which are not always adapted to European agricultural environments and farmers’ needs. The agronomic characterization of genetic resources is therefore necessary to assess new germplasm with important adaptive traits for breeding programs. One example is the INCREASE project (Bellucci et al., 2021) (https://www.pulsesincrease.eu/), which aims to characterize, at the genomic, phenotypic and phenomic levels, genetic resources representing four important legume crops in the European context: common bean (Cortinovis et al., 2021), lupin (Krock et al., 2021), chickpea (Rocchetti et al., 2022) and lentil (Guerra Garcia et al., 2021). A large set of lentil germplasm in the INCREASE project is provided by the Canadian project AGILE, which has already characterized a diverse panel of 324 accessions at the genotypic and phenotypic levels (Wright et al., 2021).

Here, we compare the agronomic potential of a set of genetic resources derived from the above-mentioned projects to local germplasm by targeting Mediterranean environments. To reach this goal, we established a diverse panel of 46 lentil genotypes composed of samples from the AGILE collection, Italian landraces and European cultivars. A 3-year trial was carried out to characterize the panel agronomically at two different localities in central and southern Italy, across two sowing seasons, allowing us to test a wide range of scenarios. Data from a subset of 16 genotypes grown in all the environments were used to dissect the genotype × environment interaction (GEI) variance. We used advanced statistical methods to evaluate the genetic potential and stability of different genotypes across various environments. We estimated breeding values (Piepho et al., 1994, 1998) and heritability (Falconer and Mackay, 1996) to identify which traits are most influenced by genetics *versus* environment, as well as multivariate analyses (AMMI and GGE) were carried out to dissect the complex interactions between genotypes and environments, helping in selection of genotypes that perform well and consistently across different conditions (Gauch, 1988; van Eeuwijk, 1995; Yan et al., 2000). Additionally, WAASB stability index (Olivoto et al., 2019) provided insights into which genotypes are most reliable and adaptable, supporting the development of resilient, high-performing varieties tailored to specific regions. Overall, these analyses are crucial for identifying genotypes well-adapted to specific environments and/or stable across different environmental conditions, making informed breeding decisions and improving crop performance in targeted environments as demonstrated by recent studies that evaluated advanced lentil genotypes (breeding lines and Iranian materials) and identified interesting materials (Shobeiri et al., 2024a, 2024b; Pezeshkpour et al., 2025).

## 2. Materials and methods

### 2.1 Plant materials

The 46 lentil genotypes evaluated in this study included 30 of the 324 accessions previously characterized agronomically during a replicated field trial carried out in 2017 in southern Italy (Metaponto: latitude 40,39401, longitude 16,78636) as part of the AGILE project (Wright et al. 2021; https://knowpulse.usask.ca/AGILE; information on the accessions and details on the field trial are provided in Wright et al. 2021). The AGILE panel mostly comprises landraces from 45 different countries. The main criteria for the selection of the 30 AGILE accessions were (i) to include genotypes with the highest yields in the 2017 field trial of the whole AGILE collection in Italy and (ii) to include genotypes originating from different geographic areas. Sixteen additional accessions were included, comprising 11 Italian landraces collected from local farmers and seed cooperatives, three Italian cultivars provided by the Società Produttori Sementi, and two French cultivars provided by Agri Obtentions (Table 1). The 16 added genotypes represent the traditional varieties and modern cultivars mostly cultivated in the areas where field trials were carried out. Single seed descent lines were used in the field trials to reduce heterogeneity within genotypes. Most of the selected genotypes were landraces from 15 different countries representing the Indian subcontinent, Mediterranean Basin and Middle East.

**Table 1.**
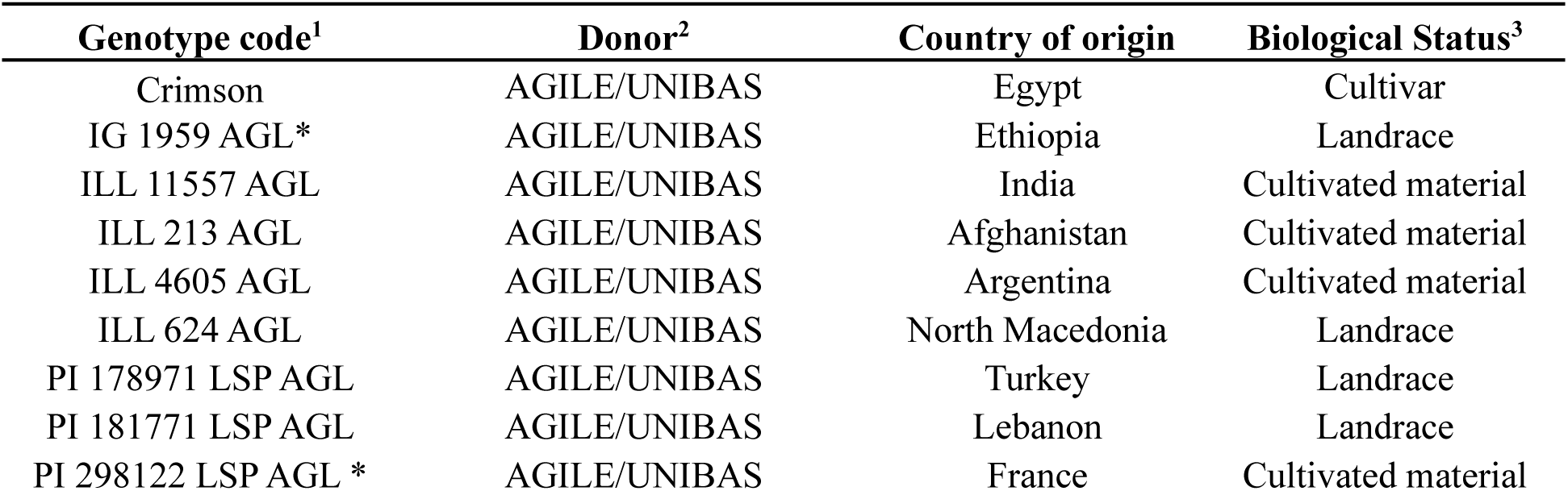
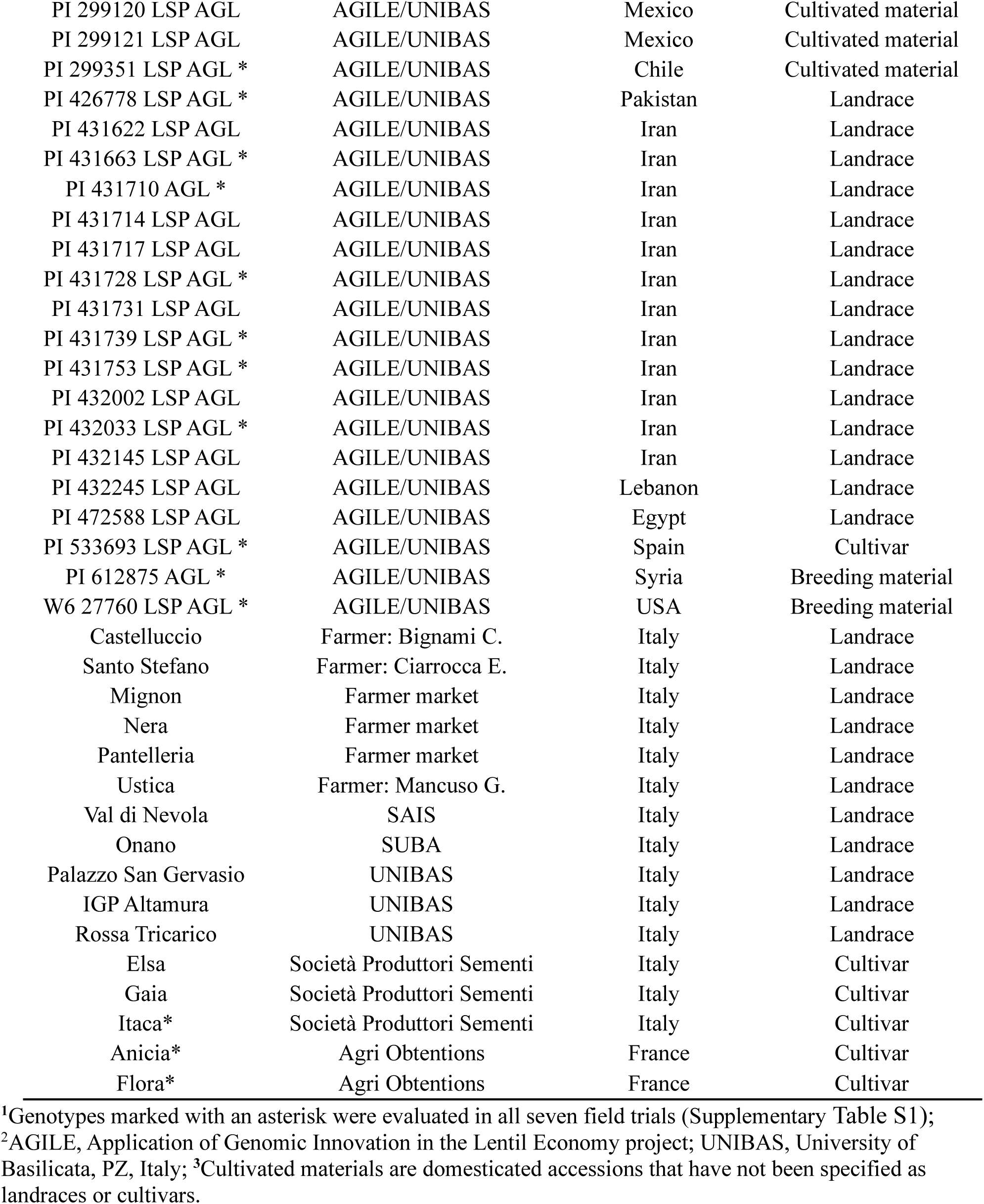
Plant materials used in this study, identified by genotype code, donor, country of origin (i.e., collection site) and biological status.

### 2.2 Multi-environmental trials

Seven agronomic trials were carried out over 3 years (2019, 2020 and 2021) and two sowing seasons (autumn and spring) in two localities (Table 2). The first was the experimental station of the Council for Agricultural Research and Economics – Cereal and Industrial Crops (CREA-CI) in Osimo, Ancona, central Italy (latitude 43.463794, longitude 13.496916). The second was the experimental station of the Agenzia Lucana di Sviluppo e di Innovazione in Agricoltura (ALSIA) in Metaponto, Potenza, southern Italy (latitude 40,39401, longitude 16,78636). We phenotypically characterized 46 genotypes (Supplementary Table S1), 16 of which (those marked with an asterisk in Table 1) were grown and successfully characterized in all seven trials.

**Table 2.** The seven field trials carried out in the present study. The trials were carried out at two localities (Osimo and Metaponto), over 3 years of cultivation (2019, 2020 and 2021) and two sowing seasons (autumn and spring).

| Experiment code<sup>1</sup> | Locality | Year | Sowing season |
| --- | --- | --- | --- |
| MA_2019 | Metaponto | 2019 | Autumn |
| MA_2020 | Metaponto | 2020 | Autumn |
| OA_2019 | Osimo | 2019 | Autumn |
| OA_2020 | Osimo | 2020 | Autumn |
| OA_2021 | Osimo | 2021 | Autumn |
| OS_2020 | Osimo | 2020 | Spring |
| OS_2021 | Osimo | 2021 | Spring |

<sup>1</sup>Experiment codes indicate the locality, year and sowing season (A = autumn [December] and S = spring [March]).

Field trials were conducted using a randomized complete block design with three replicates, in experimental plots of 10 m^2^ comprising six rows spaced 0.15 m apart. Seeds were sown in December and March for the autumn and spring seasons, respectively (Table 2), achieving a density of 130 plants/m^2^. Weed control involved pre-emergence applications of the herbicide pendimethalin. At physiological maturity, entire plants were harvested and dried in the open air. Seeds were recovered from pods using a plot harvester. Supplementary Fig. S1 shows images of OA_2021, OS_2021 and MA_2019 field trials.

Environmental data were collected in the two localities during the 3 years of trials. The following weather variables were recorded from the field weather stations: average, maximum and minimum temperatures (°C) and rainfall (mm). Thermo-pluviometric graphs for the two locations and 3 years of trials are shown in Supplementary Fig. S2.

### 2.3 Phenotypic data collection

The lentil panel was evaluated for agronomic and phenological traits relating to earliness, crop architecture and yield. Specifically, we recorded first flower (FirstF), defined as days from sowing date until 10% of plants have one open flower; first pod (FirstP), defined as days from sowing date until 10% of plants carried at least one visible pod (i.e., pods were visible without removing flower petals); plant height (PH), defined as the distance (cm) from the soil to the end of the longest stem when the plant is stretched; canopy height (CH), defined as the distance (cm) from the soil to the end of the longest stem without stretching; first pod height (FPH), defined as the distance (cm) from the soil to the first pod without stretching; yield (YLD) over the entire plot (g/plot); and seed weight (SW), defined as the 100 seed weight (g). Plant height, canopy height, and height of the first pod were recorded on three randomly-chosen plants in each plot.

### 2.4 Data analysis

Breeding value and heritability were estimated by evaluating the entire set of 46 genotypes. Assuming a linear model in Equation 1, the phenotypic value *P* for a trait and for an individual in a given environment, is the function of the overall mean (μ), the genetic effect (*G*), the environmental effect (*E*), the genotype ×environment interaction (*GEI*) and the random residual effect (*e*) within each environment, which is assumed to be normally distributed: e ∼ N(0, σ²).

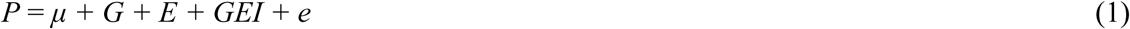

From the linearity assumption, we estimated the genotypic, environmental and GEI components of the phenotypic variance using the residual maximum likelihood procedure, considering G, E and GEI as random effects.

To predict breeding values, we modeled G and GEI as random and E as a fixed effect, enabling the estimation of best linear unbiased predictions for genotypes (BLUPg; Piepho et al., 1994, 1998). This modeling approach assumes that the sampled environments are representative of the target conditions in the central Mediterranean region, aligning with the breeding objective of selecting genotypes adapted to this specific context.

Considering that heritability can be defined as the regression of breeding values on individual phenotypes (Falconer and Mackay, 1996), we estimated the broad-sense heritability, for the unbalanced dataset of 46 genotypes, using the regression coefficient (slope of the regression) between the BLUPg and the relative phenotypic value, as shown in Equation 2.

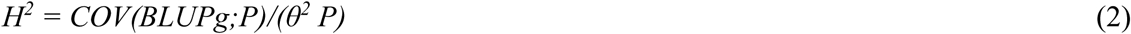

where COV(BLUPg;P) represents the covariance between the BLUP genotypic value and phenotypic value, and θ^2^ P is the total phenotypic variance. This approach implies the linearity between genetic merit and observed phenotypes and that BLUPs are unbiased predictors of true genetic values. For all traits, the predicted means of the genotypes were plotted using a caterpillar plot with horizontal bars representing the 95% confidence interval of prediction based on a two-tailed t-test (Olivoto et al., 2019).

Phenotypic data collected for the 16 genotypes in common among all trials were used to estimate trait correlation and dissect Genotypes by Environment Interaction (GEI).

Principal component analysis was run for all traits to investigate the effective agronomic space, considering estimated BLUPs for all trait-environment combinations, while Pearson correlation on BLUPs was used to determine the specific genetic correlations among traits.

Based on Equation 1, we used ANOVA to test the significance of G, E and GEI variance components considering them as fixed factors.

Furthermore, we dissected E into locality (L) and sowing season (S) effects, with the corresponding interactions: G*L (genotype × locality) and G*S (genotype × sowing season).

To analyze the structure of genotype × environment interaction (GEI) across traits, we applied the Additive Main Effects and Multiplicative Interaction (AMMI) model (Gauch, 1988; van Eeuwijk, 1995) and Genotype main effects and genotype × environment interaction effects (GGE) models (Yan et al., 2000).

AMMI is a linear model mainly used in a fixed-effect model framework. It combines two statistical procedures: ANOVA and singular value decomposition (SVD) to dissect GEI variance into different IPCAs. The resulting genotypes and environments scores are then used to construct biplots providing a graphical interpretation of the GEI effects. For the yield trait, we additionally employed the GGE model, which capture the joint effects of both the genotypic main effect G and GEI. GGE biplots provide a comprehensive visualization of genotype performance by approximating the overall effect given by G + GEI, helping identify genotypes with high performance and stability across environments.

Stability across environments was analyzed using the Weighted average absolute scores of BLUPs (WAASB) stability index (Olivoto et al., 2019), which applies SVD to the BLUP matrix for the GEI effect derived from a linear mixed model where genotypes and GEI are treated as random effects. The genotype with the lowest WAASB score is considered the most stable, deviating least from the average performance across environments. The model was applied for yield and architectural traits, and the results were displayed as a scatter plot with the abscissa represented by the performance of phenotypic trait values and the ordinate by the WAASB index. AMMI, GGE, BLUP estimates and WAASB index results were obtained using the R package (metan) (Olivoto et al., 2019) available on GitHub: https://tiagoolivoto.github.io/metan/articles/vignettes_gge.html#the-gge-model.

## 3. Results

### 3.1 Selection of plant materials

To identify plant materials adapted to Mediterranean environments, we screened yield-related data from the field trial carried out in Metaponto (southern Italy) in 2017 for the AGILE collection (Wright et al., 2020). Considering the entire set of 324 AGILE genotypes, the coefficient of variation (CV) for yield was high (62%), with an average of 114.75 g/m^2^ (Supplementary Fig. S3). We selected the 30 best-performing genotypes, covering most of the geographic areas represented in the entire AGILE collection, particularly the Indian subcontinent, Mediterranean Basin and Middle East. To make comparisons with genotypes cultivated in Italy and the Mediterranean Basin, we included 16 further genotypes representing Italian landraces and European cultivars (Table 1). We then carried out seven field trials (Table 2) using the genotypes listed in Supplementary Table S1.

### 3.2 Heritability and genotypic value

We investigated the contribution of genotypic, environmental and GEI components to the observed phenotypic variance. The proportion of genetic variance over the total phenotypic variance (i.e., heritability) for each trait is estimated in Table 3. Flowering and fruit setting times showed high heritability (H^2^ = 0.81 and 0.68, respectively) indicating a strong genetic contribution. However, environmental effect, accounted for 95% of total variance, mainly due to differences between the two sowing seasons that affected the number of days to flowering and fruit setting after sowing. Among the architectural traits, plant height, canopy height and first pod height showed intermediate to high heritability (H^2^ = 0.45, 0.50 and 0.70, respectively). For canopy height and first pod height, the residual component was high (>30%). For these traits, we also observed relatively large genotypic components (12.8% and 26.7%, respectively) and GEI effects (22.2% and 14.4%, respectively). The genotypic component of plant height was 6.67%, whereas the environmental component was 69.9% and the residual component was 15.7%, lower than that of the other architectural traits. Production traits showed relatively high heritability (H^2^ = 0.66 and 0.87 for yield and seed weight, respectively). Yield was characterized by a genotypic component of 18.39%, an environmental component of 21.77%, and the largest observed GEI effect of 35.38%. The average seed weight was characterized by a very large genotypic component (64.8%).

**Table 3.** Contribution of genotype (Gen), environment (Env), replication within the environment (rep), genotype × environment interaction (GEI), and residual components (e) to the phenotypic variation of recorded traits, along with broad sense heritability (H^2^) estimates and the coefficient of determination (R^2^).

| Phenotypic traits | Gen | Env | rep | GEI | e | $H^2$ | $R^2$ |
| --- | --- | --- | --- | --- | --- | --- | --- |
| First flower* | 3.34 | 95.40 | 0.08 | 0.89 | 0.30 | 0.81 | 0.77 |
| First pod* | 2.41 | 95.00 | 0.19 | 0.81 | 1.62 | 0.68 | 0.71 |
| Canopy height | 12.80 | 14.60 | 3.57 | 22.20 | 46.80 | 0.50 | 0.83 |
| Plant height | 6.67 | 69.90 | 1.15 | 6.56 | 15.70 | 0.45 | 0.61 |
| First pod height | 26.70 | 23.10 | 4.97 | 14.40 | 30.80 | 0.70 | 0.96 |
| Yield | 18.39 | 21.77 | 3.37 | 35.38 | 21.07 | 0.66 | 0.82 |
| Seed weight | 64.80 | 6.09 | 0.17 | 12.40 | 16.60 | 0.87 | 0.97 |

\*Nera and Pantelleria genotypes were tested only once (autumn sowing season) and were omitted from this analysis.

To estimate the genetic merit or genotypic value of the 46 lentil genotypes, we used best linear unbiased predictor (BLUP) methodology (Fig. 1; Supplementary Table S2). The lines with the tallest plants were mostly from Italy (five landraces: Palazzo San Gervaso, Onano, IGP Altamura, Pantelleria and Rossa Tricarico) and Europe (the French cultivars Flora and Anicia, and one landrace from North Macedonia), with only two landraces from Iran (PI 432002 LSP AGL and PI 432033 LSP AGL) (Fig. 1a). For the same trait, 10 lines showed BLUP confidence intervals beyond the population mean, four of which were Iranian landraces (PI 431710 LSP AGL, PI 431717 LSP AGL, PI 431753 LSP AGL, and PI 432145 LSP AGL), whereas the other six were represented by diverse materials from different geographic areas (Fig. 1a). Five genotypes had a significantly higher canopy height than the population mean, three from Italy (landrace Rossa Tricarico, and cultivars Itaca and Elsa) and two cultivated materials from Mexico, whereas the lowest canopy heights were observed for six Iranian landraces and a cultivated material from Argentina (Fig. 1a).

**Figure 1.**
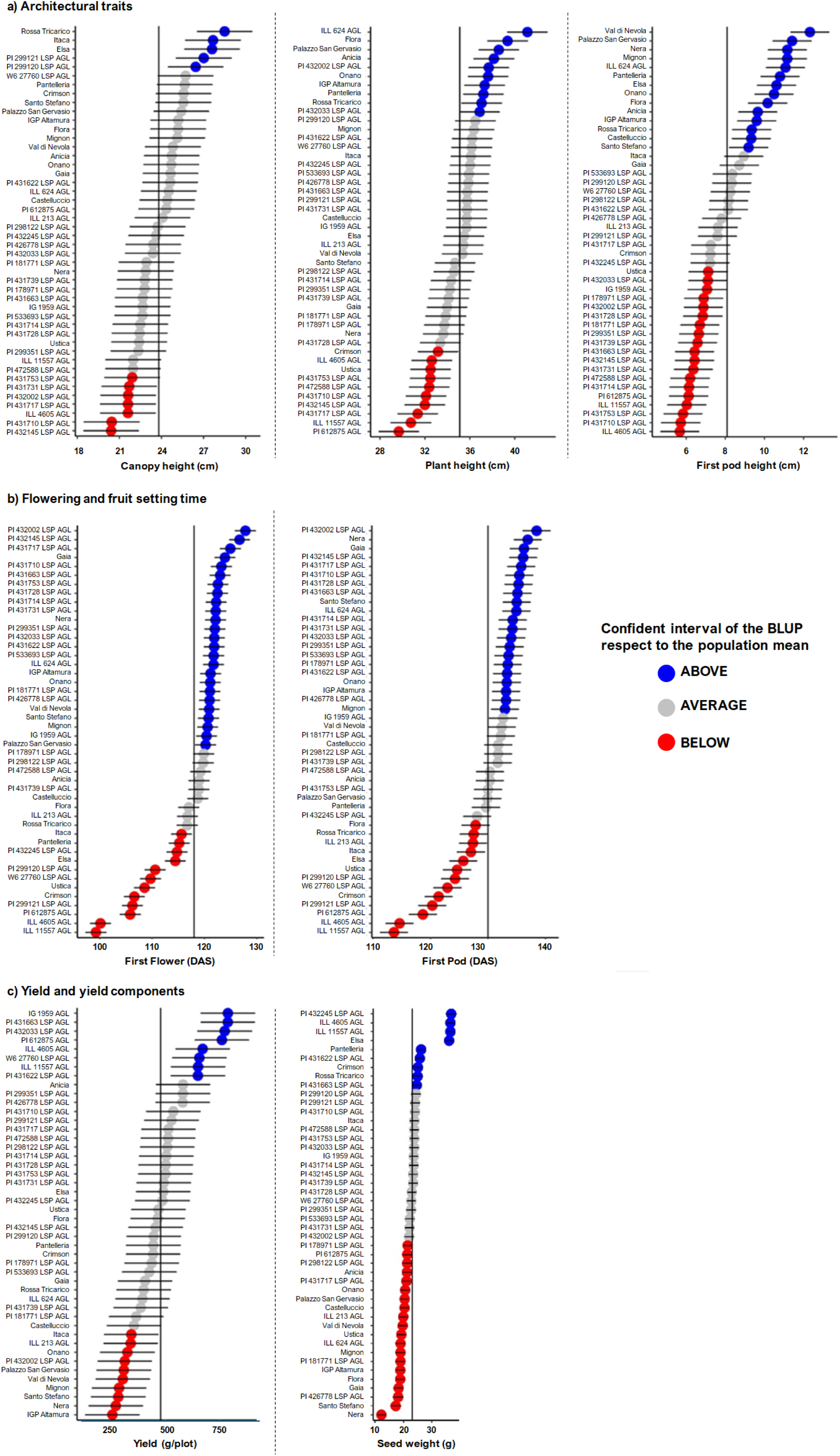
Best linear unbiased prediction (BLUP) of 46 lentil lines for **a)** architectural traits, **b)** flowering and fruit-setting times, and **c)** yield and yield components. Lines with BLUPs above and below the population mean are highlighted in blue and red, respectively. Horizontal bars represent genotype 95% confidence intervals (CI).

We also recorded the first pod height trait, which is important for the mechanical harvesting of legume crops to minimize the loss of seeds. Fourteen lines achieved the highest genotypic values for this trait, 10 of which were Italian landraces, all except one (Ustica landrace) present in the initial set of materials. The other four lines were represented by three cultivars (Elsa from Italy, and Anicia and Flora from France) and the North Macedonian landrace ILL 624 AGL (Fig. 1a). Eighteen lines, all from outside Europe, showed BLUP confidence intervals significantly lower that the population mean, the majority of which were Iranian landraces (Fig. 1a).

We observed wide variation for flowering and fruit setting time, with ranges of 28 and 25 days, respectively (Fig. 1b). The earliest flowering and fruit setting lines were ILL 11557 AGL and ILL 4605 AGL, the first from India and the second a cultivar from Argentina named Precoz. Further lines with intervals below the population means for these traits included two domesticated genotypes from Mexico, three cultivars (Elsa, Itaca and Crimson), two breeding lines (PI612875 AGL and W6 27760 LSP AGL) and an Italian landrace (Ustica) (Fig. 1b). Late-flowering genotypes (BLUP confidence intervals beyond the population mean) included all the landraces from Iran except PI431739 LSP, seven landraces from Italy, one landrace each from North Macedonia, Ethiopia, Lebanon and Pakistan, a domesticated line from Chile, and two cultivars. Most of them also showed a late fruit-setting phenotype (Fig. 1b).

Eight lines achieved the best genetic value for yield: four landraces (IG 1959 AGL from Ethiopia as well as PI 431622 LSP AGL, PI 431663 LSP AGL and PI 432033 LSP AGL from Iran), ILL 4605 AGL and ILL 11557 AGL (cultivated material from Argentina and India, respectively), and breeding lines from Syria (PI 612875 AGL) and the USA (W6 27760 AGL) (Fig. 1c). Seven Italian landraces showed the worst genetic value for yield, along with the lines Itaca, ILL 213 AGL and PI 432002 LSP AGL (Fig. 1c).

Seed weight data confirmed the distinction between *macrosperma* and *microsperma* types. The highest seed weight was observed for lines ILL 4605 AGL, PI 432245 LSP AGL, ILL 11557 AGL and Elsa (Fig. 1c). Five additional lines showed confidence intervals beyond the mean of the population, three from Italy (landraces Pantelleria and Rossa Tricarico, and cultivar Elsa), and one domesticated line each from India and Argentina. The seed weight of most Italian landraces was significantly below the population mean (Fig. 1c).

### 3.3 Phenotypic trait correlations and GEI effects

To evaluate genetic trait correlations and dissect the specific components of GEI variance component, we used the fully balanced dataset comprising 16 genotypes assessed across all seven field trials.

To obtain an overview of genotypes across the multivariate agronomic space, we performed a PCA considering the estimated BLUP for all traits based on trait–environment combinations. Principal component 1 (PC1) and PC2 explained 33.4% and 23.6% of the total variance, respectively, and showed a clear separation between genotypes based on biological status (Fig. 2). Specifically, the score plot showed that landraces, mostly from Iran, were characterized by late flowering (FirstF, FirstP) and exhibited higher productivity under spring sowing. In contrast local cultivars showed higher values for architectural traits (Fig. 2), while breeding materials were early flowering and showed greater productivity under autumn sowing (Fig. 2).

**Figure 2.**
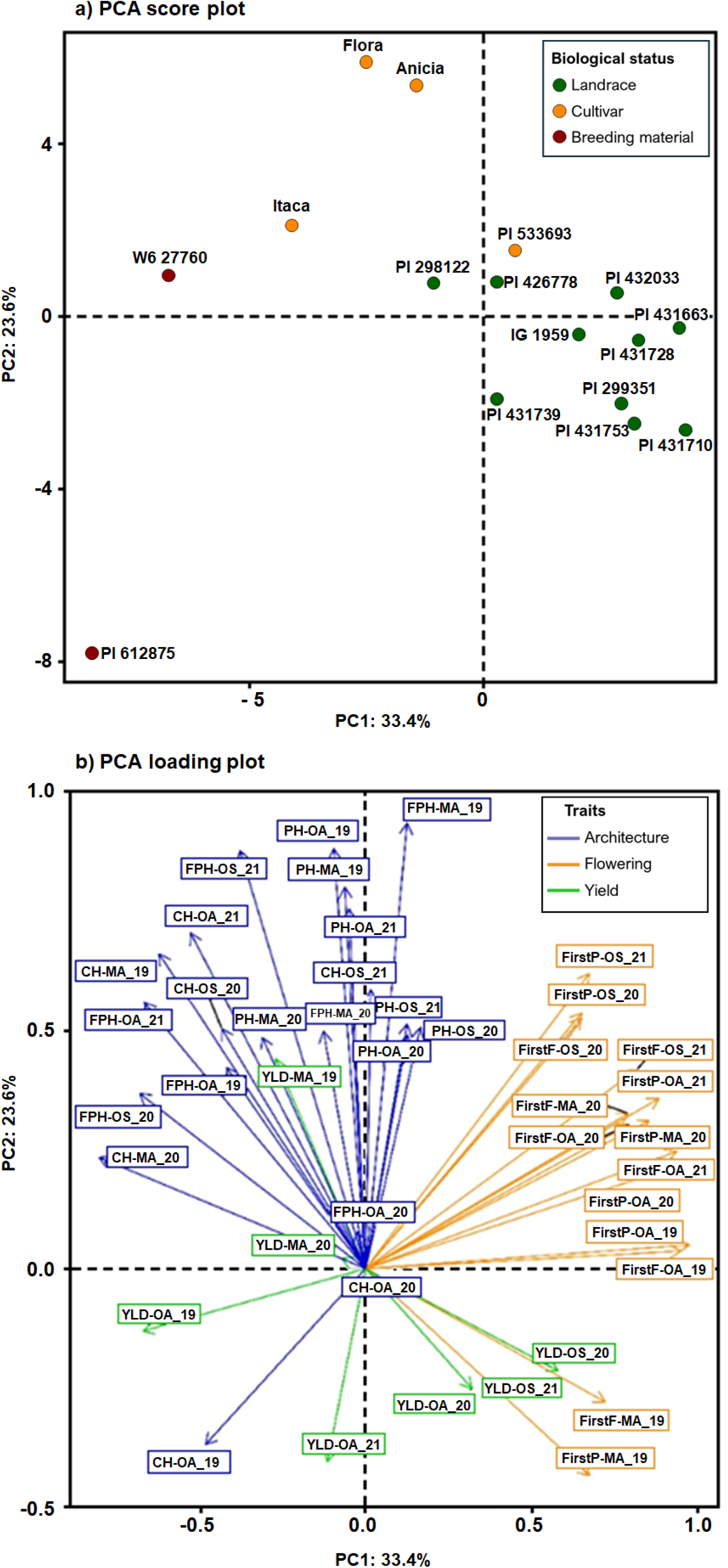
Principal component analysis carried out using the BLUPs for all trait-environment combinations. **a)** Score plot of the genotypes highlighted with different colors based on biological status: landrace (green), breeding material (red) and cultivar (orange). **b)** The loading plot for all trait– environment combinations, with flowering, architectural and yield or yield component traits highlighted in orange, blue and green, respectively.

A significant positive genetic correlation (r = 0.99; P < 0.001) was observed between flowering (FirstF) and fruit setting (FirstP), and both of these traits showed a significant negative genetic correlation with canopy height (r = –0.66 and –0.68 for canopy height/FirstF and canopy height/FirstP, respectively; P < 0.001) (Supplementary Fig. S3). First pod height was significantly positively correlated with canopy height (r = 0.70; P < 0.01) and plant height (r = 0.83 P < 0.001). Finally, seed weight showed a significant negative genetic correlation with first pod height (r = –0.59; P < 0.05) (Supplementary Fig. S4).

For architectural and production traits, significant genotype, environment and GEI effects were found by analysis of variance (ANOVA; Supplementary Table S3). The GEI effect was analyzed in detail using additive main effects and multiplicative interaction (AMMI) modeling, which allows GEI variance to be dissected into different interaction principal components (IPCs; Supplementary Table S4). Focusing on yield, a significant GEI effect was found accounting for 33.8% of total phenotypic variance with the first two IPCs explaining 79.7% of the relative GEI variance. To investigate the genotypic and environmental profiles that generate this interaction, we used AMMI biplots to show the adaptability and stability of genotypes in the tested environments (Fig. 3). In the AMMI1 and AMMI2 biplots, genotypes that cluster together are characterized by similar performance, and environments represented by vectors that form an acute angle are positively correlated.

**Figure 3.**
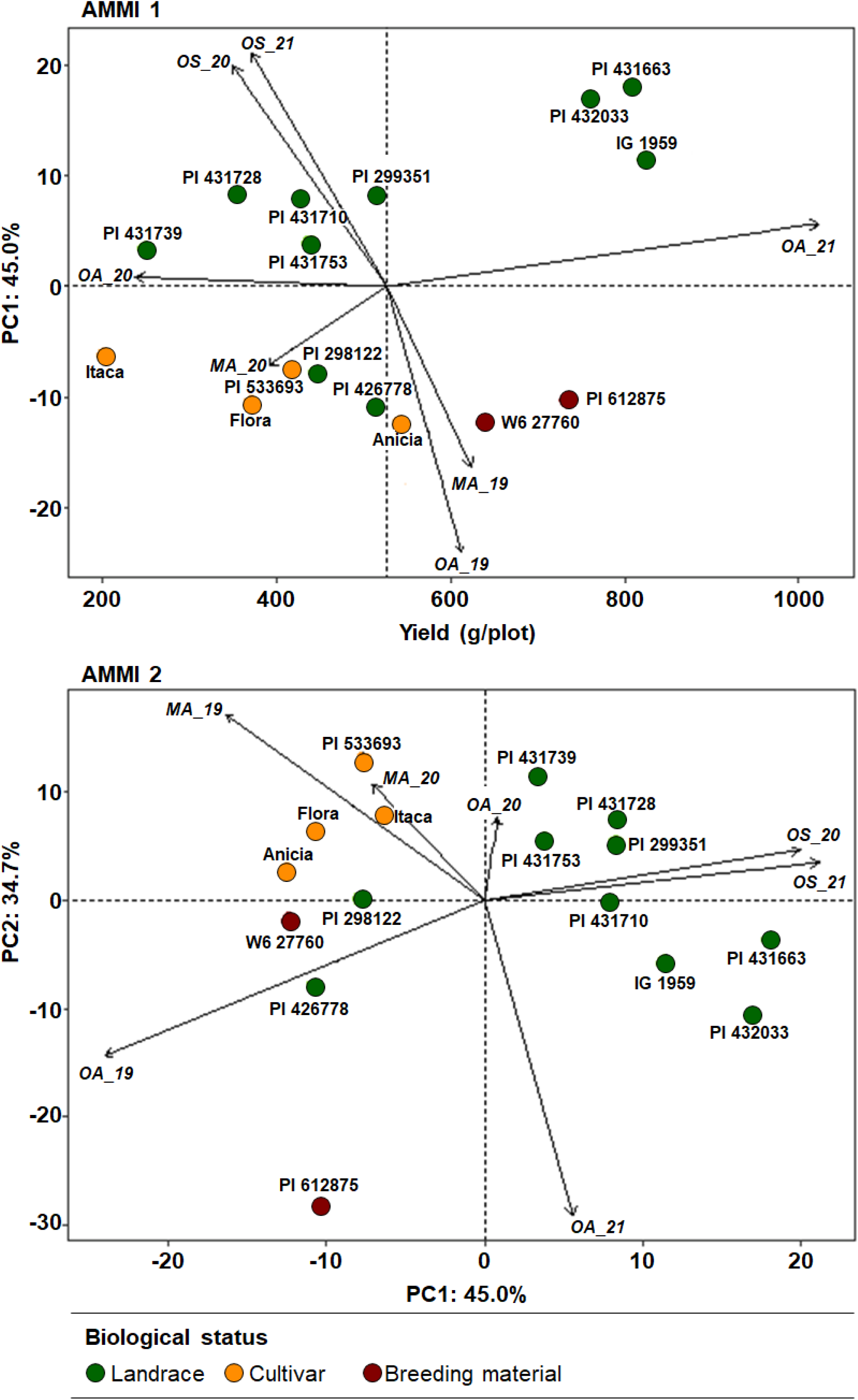
AMMI biplots for yield data from 16 lentil genotypes grown in seven environments based on genotypic and environmental scores. **a)** AMMI1 biplot in the abscissa the genotype and environmental yield and in the ordinates the PC1 scores. **b)** AMMI2 biplot (PC1 vs PC2) for plot yield data. Dots represent the 16 genotypes while the arrows the seven different environments.

The AMMI1 biplot represents the IPC score against the mean grain yield for all the environments and genotypes (Fig. 3). IPC1 retained 45% of the total GEI variance and mainly separated the spring and autumn sowing seasons, with landraces tending to perform better in the spring whereas cultivars and breeding lines performed better in the autumn. The breeding lines PI 612875 AGL and W6 27760 LSP AGL performed better in the autumn sowing seasons, with yield exceeding the population average, whereas landraces PI 4316163 LSP AGL, PI 432033 LSP AGL and IG 1959 AGL achieved a higher yield in the spring sowing seasons. This reflects a differential response to the primary environmental contrasts between autumn and spring sowing. Specifically, autumn sowing is characterized by slower thermal accumulation during the vegetative phase (mean daily temperature ∼7.5°C) under short-day photoperiods, whereas spring sowing is associated with more rapid thermal accumulation (mean daily temperature ∼15°C) combined with long-day photoperiods (Fig.S1).

In AMMI2, the scores for IPC1 and IPC2 explained 45.0% and 34.7% of the total GEI variance, respectively (Fig. 3). IPC1 separated environments according to the two sowing seasons, whereas IPC2 tended to separate environments according to localities (Osimo *vs* Metaponto). As for AMMI1, the AMMI2 results confirmed a positive interaction between landraces and the spring sowing season, whereas cultivars performed best in the Metaponto locality. Strong positive interaction was also observed for the breeding lines P612875 in Osimo autumn trials, particularly Osimo_2019_A and Osimo_2021_A.

To determine the total genetic variation attributable to the joint effect of genotype and GEI, we applied genotype plus genotype–environment interaction (GGE) analysis (Yan et al., 2000). The so-called “which-won-where” view of the GGE biplot is shown for yield in Fig. 4. The partitioning of GEI in the GGE biplot showed that PC1 and PC2 accounted for 54.6% and 27.7% of the GGE sum of squares, respectively, explaining 82.3% of the total variation. Seven environments, given by the combination of locations and sowing seasons, fell into two distinct sectors, one including the autumn trials carried out in both Osimo and Metaponto (Osimo_2019_A, Osimo_2021_A, Metaponto_2019_A and Metaponto_2020_A) and the other mainly the spring trials (Osimo_2020_S and Osimo_2021_S), along with Osimo_2020_A (Fig. 4). The winning genotypes in the sector containing the autumn sowing season environments were the breeding lines PI612875 AGL followed by W6 27760 LSP AGL, whereas the winning genotypes in the sector containing the spring sowing season trials were the Iranian landraces PI 431663 LSP AGL, followed by PI 432033 LSP AGL and IG 1959 AGL. These results agreed with the AMMI analysis. In the section characterized by the presence of three out of four cultivars (Itaca, Flora and PI 533693), any test environment was present, suggesting that these cultivars were less productive in all the environments we considered.

**Figure 4.**
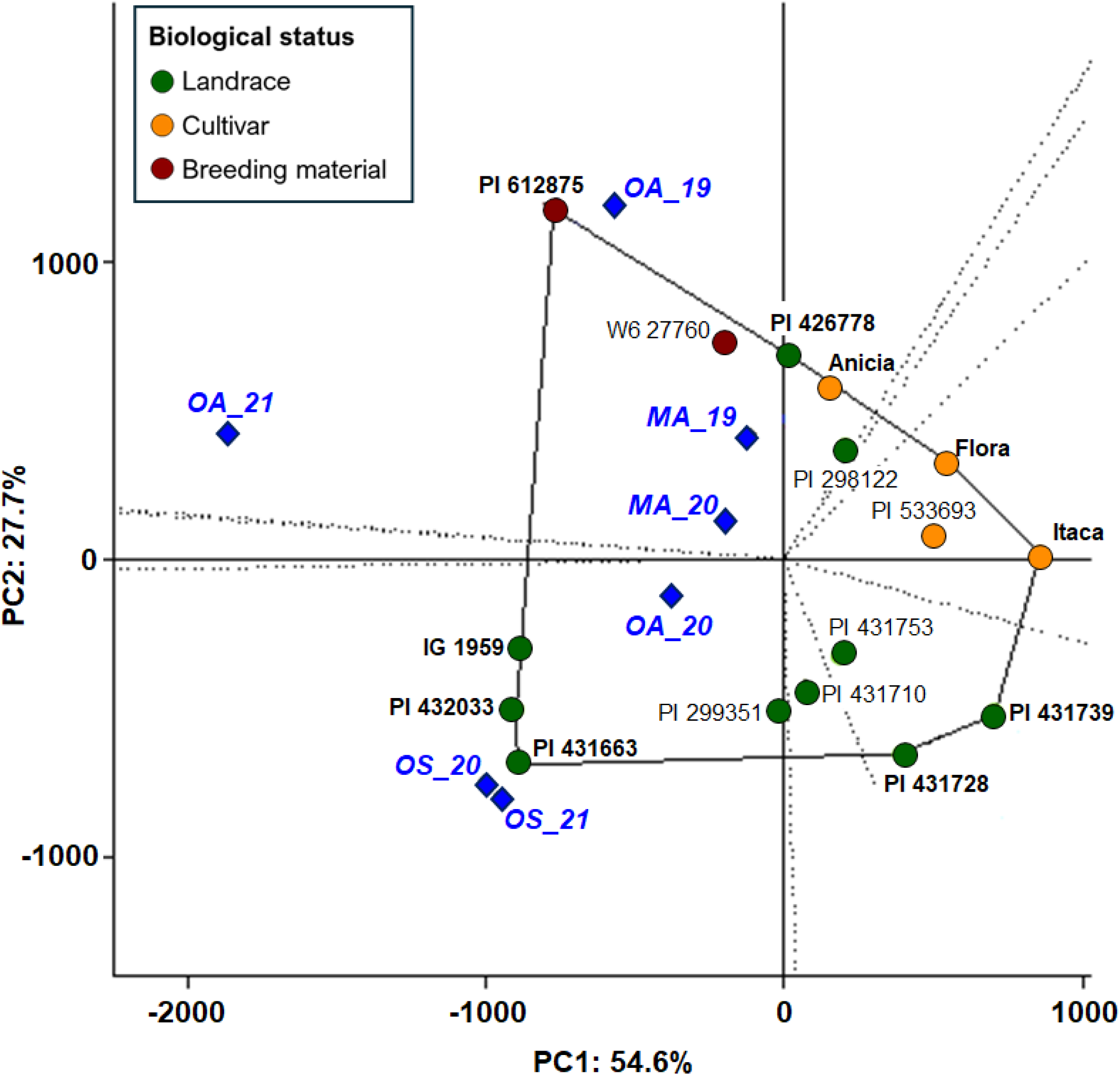
“Which-won-where” GGE biplot based on yield data for 16 genotypes. Circles are colored based on biological status in seven environments (blue diamonds). Vertex genotypes are connected with solid lines and the remaining genotypes fall within the polygon. A set of lines starting from the origin intersects each side of the polygon. The perpendicular lines to the polygon sides divide the biplot into sectors where environments that share similar ranks of the genotypes within a specific sector are grouped together and the winning genotype for a sector is the vertex genotype.

To investigate genotypic stability across environments, we used the weighted average of absolute scores (WAASB) stability index. The scatter plot with yield in the abscissa and the WAASB stability index in the ordinate allowed us to derive a joint interpretation between yield performance and stability (Fig. 5). Genotypes and environments fall in four different quadrants. For breeding purposes, the focus is the second (II) and fourth (IV) quadrants, because they group genotypes with yields above the population average. The second quadrant (II) represents highly productive but unstable genotypes, characterized by strong performance in the environments included in the quadrant, whereas the fourth quadrant (IV) contains genotypes with yields above the total average that are broadly adapted (low WAASB index) to all the tested environments.

**Figure 5.**
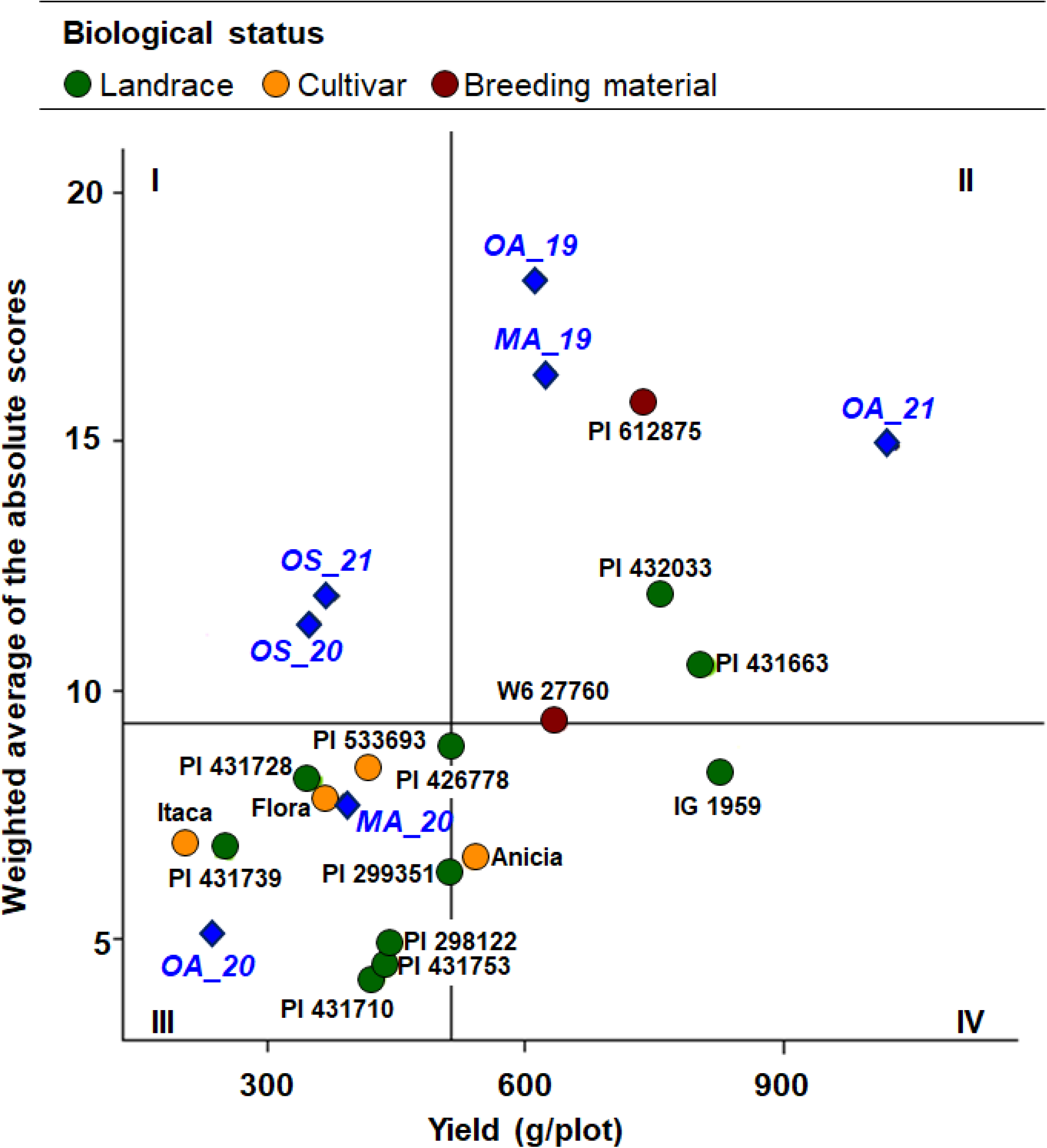
Score plot of grain yield vs weighted average of absolute scores for the best linear unbiased predictions of the genotype–environment interaction (WAASB). Genotypes are represented by colored circles based on biological status, whereas blue diamonds indicate the seven different environments. Horizontal and vertical black lines indicate the average stability and population yield, respectively.

Among the 16 genotypes characterized in all the seven environments, five showed the highest yield, as previously suggested by BLUPs estimates (Figure 1). Three of them (PI 432033 LSP AGL, PI 431663 LSP AGL and PI 612875 AGL) fell within the second (II) quadrant along with the autumn sowing trials carried out in 2019 in Metaponto and Osimo and in 2021 in Osimo. They were characterized by the highest WAASB scores, indicating they are not stable in all the environments but more adapted to autumn sowing seasons, especially PI 612875 AGL and PI 432033 LSP AGL (Figure 5). The W6 27760 LSP AGL genotype fell between quadrants II and IV and the IG 1959 AGL genotype fell within quadrant IV. Among genotypes with a higher yield than the population average, IG 1959 AGL showed the lowest WAASB score meaning a good and stable performance across all different environments (Fig. 5).

Finally, we partitioned the environmental (E) effect by distinguishing the contributions of location (L) and sowing season (S) using an ANOVA-based linear model (Supplementary Table S5).We found that lines IG 1959 AGL and PI 431663 LSP AGL were the best performing and most stable genotypes across locations and sowing seasons (Fig. 6). Comparing sowing seasons and locations, we found that the yield was significantly higher in autumn trials and the Osimo locality compared to spring sowing and Metaponto, respectively. We also found a significant genotype interaction with locality effect particularly for the PI 612875 AGL breeding line, which achieved a significantly higher yield in Osimo than in Metaponto. Considering the differences between sowing seasons, almost all the genotypes showed a significantly higher yield in autumn compared to spring sowing trials, particularly the breeding lines PI 612875 AGL and W6 27760 LSP AGL, whereas almost all of the Iranian landraces achieved a stable performance across both sowing seasons.

**Figure 6.**
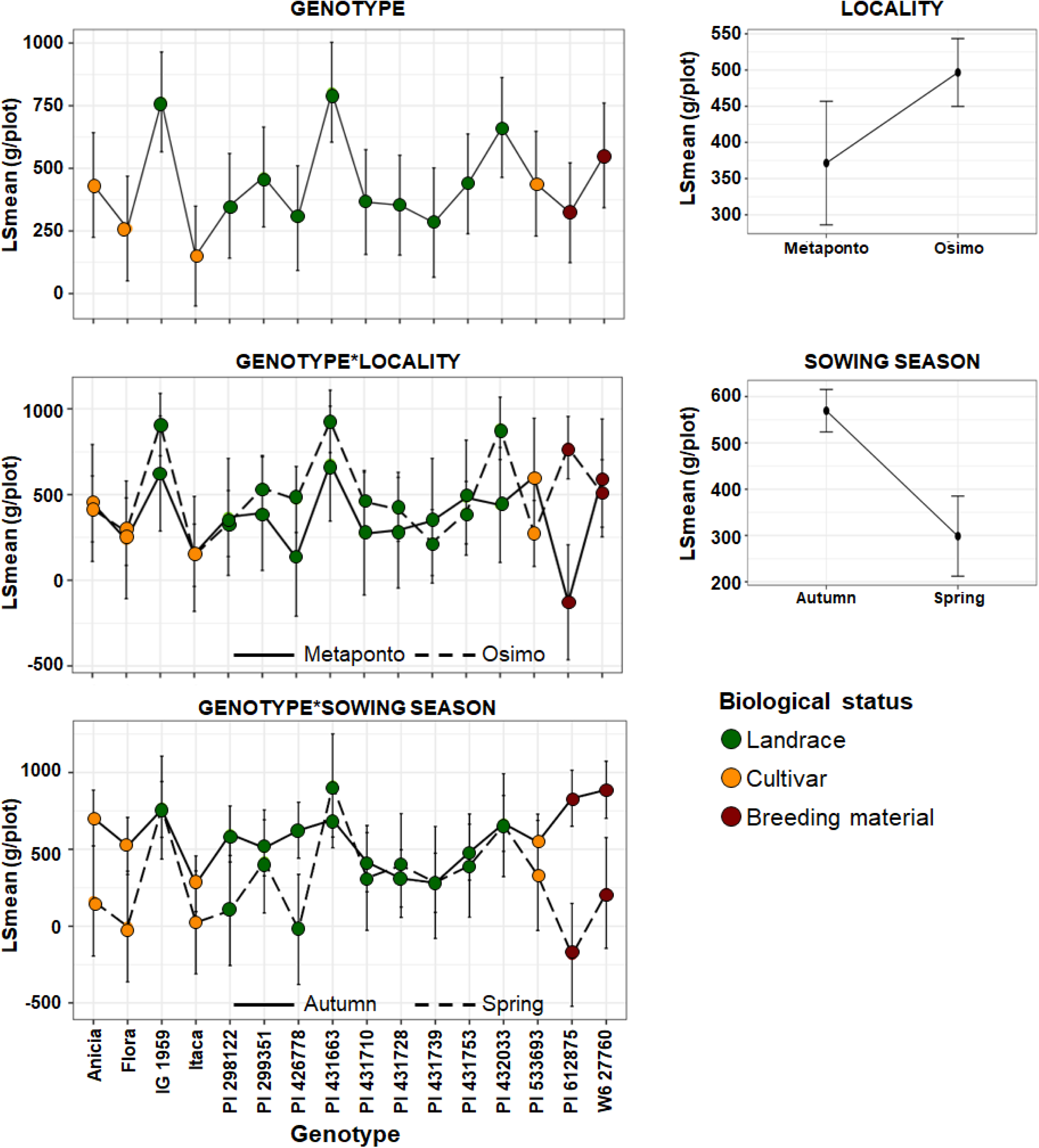
Linear model with genotype, location and season effects and interactions for the yield trait.

## 4. Discussion

Lentil is one of the oldest cultivated grain legumes and is strongly linked to European and Mediterranean food cultures. This species can therefore play a key role in supporting the transition toward a largely plant-based diet for European consumers. However, little effort has been dedicated to lentil breeding in Europe, and few lentil cultivars have been released. Research on the characterization of diverse lentil genetic resources in European and Mediterranean agro-ecosystems is also limited, with examples mainly involving the evaluation of local germplasm (Lázaro et al., 2001; Bacchi et al., 2010; Zaccardelli et al., 2012; Cristóbal et al., 2014; Ruisi et al., 2015; Gaad et al., 2018; Sellami et al., 2021; Baxevanos et al., 2024). There is increasing interest in the development of legume varieties adapted to European and Mediterranean environments, leading to funding for the characterization of diversity in legume collections, as seen in the INCREASE project (Bellucci et al., 2021; https://www.pulsesincrease.eu/), and initial publications testing a wide set of materials in Mediterranean environments (Preiti et al., 2024; Zeroual et al., 2024).

We set out to develop a diverse set of lentil materials for agronomic characterization and GEI analysis in Italian Mediterranean environments and to identify adapted germplasm for breeding purposes. We exploited an earlier field trial carried out in the south of Italy (Metaponto, Basilicata region) based on the characterization of a wide and geographically diverse panel of 324 genotypes developed in the AGILE project (Wright et al., 2021). To target new germplasm potentially adapted to Mediterranean environments, we selected the 30 best-yielding AGILE genotypes. This allowed us to maintain a geographically diverse set of genotypes, including lines based on materials originally from different geographic areas such as the Mediterranean Basin, Middle East, central Asia, the Indian subcontinent and Ethiopia. We added another 16 genotypes representing local Italian landraces and varieties that are mainly grown by farmers in Italy to ultimately compare the AGILE resources with materials already cultivated in the Italian Mediterranean region.

Phenotypic characterization, across seven environments derived by combinations of localities, sowing seasons and years, allowed us to dissect the phenotypic variance of agronomic traits in terms of genotypic, environmental and GEI components. The large collection of 46 genotypes allowed to evaluate a large portion of the genetic variance, however it was not equally replicated among the seven environments, making the use of the standard ratio of variances: Ɵ^2^_G_/(Ɵ^2^_G_ + Ɵ^2^_GEI_ + Ɵ^2^_e_) inappropriate. Instead, we calculated the regression of genotypic values on phenotypic values using genotypic BLUPs (estimates of specific genotypic values), to estimate broad-sense heritability as previously described (Falconer and Mackay, 1996).

Flowering time is an important adaptive trait and is therefore of significant interest for breeding. Varieties that mature faster could reduce yield losses due to water and heat stress during the reproductive and grain-filling stages (Naik et al., 2024). Considering that lentil is a long-day species with different photoperiod sensitivity across genotypes, we did not only test the material in different locations, but we also targeted two different sowing seasons: autumn, characterized by short-day conditions in the early phases and long-day conditions in the reproductive phases, and spring, characterized by long-day conditions since the early phases. For this reason, we detected a strong environmental component for flowering and fruit-setting time, mainly reflecting day length differences between the two sowing seasons (Summerfield et al., 1985). By estimating the genetic merit of the different genotypes, we identified those potentially superior for earliness, such as ILL 11557 AGL and ILL 4605 AGL, which are domesticated materials from India and Argentina, respectively. Among the 16 genotypes with complete datasets for all seven environments, the breeding lines PI 612875 AGL and W6 27760 LSP AGL showed the shortest cycles for flowering and fruit setting (Figure 2).

Architectural traits are important because they affect plant management and yield (Holland, 2007; Teichmann and Muhr, 2015) as well as disease control by regulating the extent of contact with pathogens, or by creating barriers or unfavorable microclimates for pathogen growth and development (Ando et al., 2007; Costes et al., 2013). Several studies have pointed out the importance of such traits in lentil, especially to improve harvesting and reduce seed losses (Erskine and Goodrich,1991; Kuzbakova et al., 2022) but also by enhancing yields (Silva-Perez et al., 2022). Phenotypic variance partitioning differed for the three architectural traits. The residual component was lowest for plant height (15%), but higher for canopy height and first pod height, probably because these latter traits are more influenced by lodging, which can create statistical noise that increases the residual component. Canopy height and first pod height also showed a relatively high genotypic effect. From genetic merit estimates, our results indicated that European materials (Italian cultivars/landraces and French cultivars) achieved the best results. Almost all the Italian landraces, with the exception of the Ustica landrace, achieved the highest values for the first pod height, which is valuable for mechanical harvesting and reducing seed loss. We observed a relatively high heritability for yield (66%) indicating an overall potential for selection at the population level for adaptation in Mediterranean environments. The best-yielding genotypes were IG 1959, PI 612875, PI 431663 LSP and PI 432033 LSP, based on landraces and breeding material originating from Ethiopia, Syria, and Iran. We did not find any local germplasm with better performance. Similar experiments of multi-location field trials conducted in Iran confirms that improved lentil cultivars developed from material originated mainly from these geographic regions can combine high yield with stability across diverse, rainfed sites (Shobeiri et al., 2024a, 2024b; Pezeshkpour et al., 2025).

Among the overall population of 46 genotypes, a subset of 16 lines (Table 3) was fully replicated across all tested environments. This common set included AGILE germplasm, comprising landraces and breeding materials capable of producing sufficient seed for all trials, as well as European cultivars used as controls. This subset enabled a detailed analysis of phenotypic trait correlations and allowed for the dissection of genotype-by-environment interaction (GEI) effects.

Principal Component Analysis (PCA), performed using all trait-by-environment combinations, revealed clear agronomic differentiation between the local European cultivars and the AGILE germplasm. While European cultivars clustered together based on higher values for architectural traits, the breeding materials and landraces displayed distinct agronomic patterns, particularly in terms of yield performance across both sowing seasons. These results indicate that the target of the former breeding efforts was to improve architectural traits, thus facilitating mechanical harvesting but also increasing yield (Silva-Perez et al., 2022).

Analyzing in detail the specific environmental effect we observed overall higher yield in autumn compared to spring sowing trials and in central rather than southern Italy. In central Italy, lentils are usually sown in spring (February to March), whereas we found that, due to rising winter temperatures and unpredictable spring drought in this area, higher yields can be achieved by sowing lentil in autumn. This was previously reported in the Mediterranean-type environments of southern Australia, where early sowing allowed a longer period of vegetative and reproductive growth, faster canopy development, more absorption of photosynthetically active radiation, higher seed yields, and better water use efficiency (Siddique et al., 1998).

Considering the relatively mild temperatures typical of the local autumn-winter growing season (7-8°C) and their projected increase under future climate scenarios (Khodayar Pardo et al., 2023), along with initially short-day photoperiods during the vegetative stage, this sowing window is likely to favor early-flowering genotypes. Due to their photoperiod sensitivity, these genotypes can complete vegetative development and initiate flowering before the onset of potential spring drought stress.

The GEI structure of yield was analyzed using AMMI and related GGE models, revealing that the main driver of GEI is the environmental difference between the autumn and spring sowing seasons. In autumn sowing, the best performing lines were the breeding material PI 612875 and W6 27760 LSP that were also included among the significant early group of genotypes. These two genotypes are differentiated for architectural traits, with PI 612875 AGL featuring a significantly lower plant height and first pod height. In this case, intercropping with cereals such as wheat or barley may be a useful strategy to increase the first pod height and facilitate stand harvesting (Carr et al., 1995).

On the other hand, we found two landraces from Iran PI 431663 and PI 432033 LSP and one from Ethiopia IG 1959, that beside well performing in autumn sowing, also had outstanding production in spring sowing, where higher temperatures and reduced rainfall occurred during the growing season We investigated the stability of the genotypes by estimating the WAASB index, which considers all IPCAs derived from the BLUP GEI variance decomposition, allowing an unbiased interpretation of genotype stability across environments (Olivotto et al., 2019). We found that the Ethiopian line IG 1959 AGL was the most stable among the best-yielding genotypes. This line represents a valuable genetic resource for the development of improved cultivars adapted to Mediterranean environments. It holds potential as a broadly adapted parental line in breeding programs.

The identified materials can be valuable sources of new diversity and can be crossed with cultivars already grown in the same areas, such as for example the Itaca variety, to combine desirable traits and address specific agronomic, farmer, and market demands.

## 5. Conclusions

Here, we exploited the availability of a wide and diverse lentil panel representing the worldwide geographic distribution of the species to select a set of materials targeting high-production and adaptation to Mediterranean agro-environments. Agronomic analysis in multiple environments revealed promising new genotypes suitable for adaptation to the Mediterranean region, and GEI analysis identified genotypes adapted either to autumn or spring sowing seasons as well as lines stable across different environmental conditions. These genotypes can be used in breeding to develop new varieties able to promote the cultivation of legumes in Europe and to contribute to sustainable development of agriculture.

## CRediT authorship contribution statement

Lorenzo Rocchetti: Data curation, Formal analysis, Investigation, Methodology, Software, Validation, Visualization, Writing - original draft. Alex Kumi Frimpong: Data curation, Formal analysis, Visualization, Writing – review and editing. Valerio Di Vittori: Formal analysis, Visualization, Writing – review and editing. Chiara Santamarina: Investigation, Formal Analysis, Writing – review and editing. Alice Pieri: Formal analysis, Visualization, Writing – review and editing. Andrea Tosoroni: Investigation, Writing – review and editing. Simone Papalini: Investigation, Writing – review and editing. Francesca Francioni: Investigation, Writing – review and editing. Evan Musari: Investigation, Writing – review and editing. Elisa Bellucci: Investigation, Writing – review and editing. Laura Nanni: Investigation, Writing – review and editing. Stefania Marzario: Investigation, Resources. Giuseppina Logozzo: Investigation, Resources. Tania Gioia: Investigation, Resources. Kirstin Bett: Resources, Supervision, Writing – review and editing. Guido Arlotti: Funding acquisition, Project Administration, Supervision; Marco Silvestri: Funding acquisition, Project Administration, Supervision; Roberto Papa: Conceptualization, Formal analysis, Funding acquisition, Methodology, Project Administration, Supervision, Writing-original draft. Elena Bitocchi: Conceptualization, Data curation, Formal analysis, Investigation, Methodology, Project Administration, Resources, Supervision, Validation, Visualization, Writing-original draft.

## Code availability

https://github.com/LorePlant/Lentil_GEI

## Data Availability

Data will be made available on request.

## References

Aguilar, O.M.; Collavino, M.M.; Mancini, U. Nodulation competitiveness and diversification of symbiosis genes in common beans from the American centers of domestication. Sci. Rep. 2022, 12, 4591. 10.1038/s41598-022-08720-0.

Alo, F.; Furman, B.J.; Akhunov, E.; Dvorak, J.; Gepts, P. Leveraging genomic resources of model species for the assessment of diversity and phylogeny in wild and domesticated lentil. J. Hered. 2011, 102, 315–329. 10.1093/jhered/esr015.

Ando, K.; Grumet, R.; Terpstra, K.; Kelly, J.D. Manipulation of plant architecture to enhance crop disease control. CAB Rev. Perspect. Agric. Vet. Sci. Nutr. Nat. Resour. 2007, 2, 1–8. 10.1079/PAVSNNR2007202.

Bacchi, M.; Leone, M.; Mercati, F.; Preiti, G.; Sunseri, F.; Monti, M. Agronomic evaluation and genetic characterization of different accessions in lentil (*Lens culinaris* Medik.). Ital. J. Agron. 2010, 5, 303–314. 10.4081/ija.2010.303.

Baxevanos, D.; Kargiotidou, A.; Noulas, C.; Kouderi, A.-M.; Aggelakoudi, M.; Petsoulas, C.; Tigka, E.; Mavromatis, A.; Tokatlidis, I.; Beslemes, D.; Vlachostergios D.N. Lentil cultivar evaluation in diverse organic Mediterranean environments. Agronomy 2024, 14, 790. 10.3390/agronomy14040790.

Bellucci, E.; Mario Aguilar, O.; Alseekh, S.; Bett, K.; Brezeanu, C.; Cook, D.; De la Rosa, L.; Delledonne, M.; Dostatny, D.F.; Ferreira, J.J.; Geffroy, V.; Ghitarrini, S.; Kroc, M.; Kumar, S.; Logozzo, G.; Marino, M.; Mary-Huard, T.; McClean, P.; Meglič, V.; Messer, T.; Muel, F.; Nanni, L.; Neumann, K.; Servalli, F.; Străjeru, S.; Varshney, R.K.; Vasconcelos, M.W.; Zaccardelli, M.; Zavarzin, A.; Bitocchi, E.; Frontoni, E.; Fernie, A.R.; Gioia, T.; Graner, A.; Guasch, L.; Prochnow, L.; Opperman, M.; Susek, K.; Tenaillon, M.; Papa, R. The increase project: Intelligent collections of food-legume genetic resources for european agrofood systems. Plant J. 2021, 108, 646–660. 10.1111/tpj.15472.

Carr, P.M.; Gardner, J.C.; Schatz, B.G.; Zwinger, S.W.; Guldan, S.J. Grain yield and weed biomass of a wheat-lentil intercrop. Agron. J. 1995, 87, 574–579. 10.2134/agronj1995.00021962008700030030x

Costes, E.; Lauri, P.E.; Simon, S.; Andrieu, B. Plant architecture, its diversity and manipulation in agronomic conditions, in relation with pest and pathogen attacks. Eur. J. Plant Pathol. 2013, 135, 455–470. 10.1007/s10658-012-0158-3.

Cortinovis, G.; Oppermann, M.; Neumann, K.; Graner, A.; Gioia, T.; Marsella, M.; Alseekh, S.; Fernie, A.R.; Papa, R.; Bellucci, E.; Bitocchi, E. Towards the development, maintenance, and standardized phenotypic characterization of Single-Seed-Descent genetic resources for common Bean. Curr Protoc. 2021, 5, e133. 10.1002/cpz1.133.

Cristóbal, M.D.; Pando, V.; Herrero, B. Morphological Characterization of lentil (*Lens culinaris* Medik.) landraces from Castilla y León, Spain. Pak. J. Bot. 2014, 46, 1373–1380.

Dabin, Z.; Pengwei, Y.; Na, Z.; Changwei, Y.; Weidong, C.; Yajun, G. Contribution of green manure legumes to nitrogen dynamics in traditional winter wheat cropping system in the Loess Plateau of China. Eur. J. Agron. 2016, 72, 47–55. 10.1016/j.eja.2015.09.012.

Erskine, W.; Goodrich, W.J. Variability in lentil growth habit. Crop Sci., 1991, 31, 1040–1044. 10.2135/cropsci1991.0011183X003100040039x.

Falconer, D.; Mackay, T. Introduction to Quantitative Genetics, 4th ed. Prentice Hall: Harlow, UK, 1996.

Gaad, D.; Laouar, M.; Abdelguerfi, A.; Gaboun, F. Collection and agro morphological characterization of Algerian accessions of lentil (*Lens culinaris*). Biodiversitas, 2018, 19, 183–193. doi: 10.13057/biodiv/d190125

Gauch, H.G.; Zobel, R.W. Predictive and postdictive success of statistical analyses of yield trials. Theor. Appl. Genet. 1988, 76, 1–10. 10.1007/BF00288824.

Gerten, D.; Heck, V.; Jägermeyr, J.; Bodirsky, B.L.; Fetzer, I.; Jalava, M.; Kummu, M.; Lucht, W.; Rockström, J.; Schaphoff, S.; Schellnhuber, H.J. Feeding ten billion people is possible within four terrestrial planetary boundaries. Nat. Sustain. 2020, 3, 200–208. 10.1038/s41893-019-0465-1.

Guerra-García, A.; Gioia, T.; von Wettberg, E.; Logozzo, G.; Papa, R.; Bitocchi, E.; Bett, K.E. Intelligent characterization of lentil genetic resources: Evolutionary history, genetic diversity of germplasm, and the need for well-represented collections. Curr. Protoc. 2021, 1, e134. 10.1002/cpz1.134.

Holland, J.B. Genetic architecture of complex traits in plants. Curr. Opin. Plant. Biol. 2007, 10, 156–161. 10.1016/j.pbi.2007.01.003.

Khodayar Pardo, S. K.; Pastor, P.; Valiente, J.A.; Paredes-Fortuny, L.; Benetó, P. The new reality of the Mediterranean: accelerating impacts of climate change EGU General Assembly 2023, Vienna, Austria, 24–28 Apr 2023, EGU23-15233, 2023. 10.5194/egusphere-egu23-15233.

Kroc, M.; Tomaszewska, M.; Czepiel, K.; Bitocchi, E.; Oppermann, M.; Neumann, K.; Guasch, L.; Bellucci, E.; Alseekh, S.; Graner, A.; Fernie, A.R.; Papa, R.; Susek, K. Towards Development, Maintenance, and Standardized Phenotypic Characterization of Single-Seed-Descent Genetic Resources for Lupins. Curr. Protoc. 2021, 1, e191. 10.1002/cpz1.191.

Kuzbakova, M.; Khassanova, G.; Oshergina, I.; Ten, E.; Jatayev, S.; Yerzhebayeva, R.; Bulatova, K.; Khalbayeva, S.; Schramm, C.; Anderson, P.; Sweetman, S.; Jenkins, C.L.D.; Soole, K.L.; Shavrukov, Y. Height to First Pod: A review of genetic and breeding approaches to improve combine harvesting in legume crops. Front. Plant Sci. 2022, 13, 948099. 10.3389/fpls.2022.948099.

Ladizinsky, G. The origin of lentil and its wild genepool. Euphytica 1979, 28, 179–187. 10.1007/BF00029189.

Laghetti, G.; Piergiovanni, A.R.; Sonnante, G.; Lioi, L.; Pignone, D. The Italian lentil genetic resources: a worthy basic tool for breeders. Europ. J. Plant Sci. Biotech. 2008, 2, 48–59.

Lázaro, A.; Ruiz, M.; de la Rosa, L.; Martín, I. Relationships between agro/morphological characters and climatic parameters in Spanish landraces of lentil (*Lens culinaris* Medik.). Genet. Resour. Crop. Evol. 2001, 48, 239–249. 10.1023/A:1011234126154.

Lindström, K.; Mousavi, S.A. Effectiveness of nitrogen fixation in rhizobia. Microb. Biotechnol. 2020, 13, 1314–1335. 10.1111/1751-7915.13517.

Naik, Y.D.; Sharma, V.K.; Aski, M.S.; Rangari, S.K.; Kumar, R.; Dikshit, H.K.; Sahani, S.; Kant, R.; Mishra, G.; Mir, R.R.; Kudapa, H. Phenotypic profiling of lentil (*Lens culinaris* Medikus) accessions enabled identification of promising lines for use in breeding for high yield, early flowering and desirable traits. Plant Gen. Resour.: Characterisation Util., 2024, 22, 69–77. 10.1017/S1479262124000042.

Nartea, A.; Kuhalskaya, A.; Fanesi, B.; Orhotohwo, O.L.; Susek, K.; Rocchetti, L.; Di Vittori, V.; Bitocchi, E.; Pacetti, D.; Papa, R. Legume byproducts as ingredients for food applications: Preparation, nutrition, bioactivity, and techno-functional properties. Compr. Rev. Food Sci. Food Saf. 2023, 22, 1953–1985. 10.1111/1541-4337.13137.

Olivoto, T.; Lúcio, A.D.C.; da Silva, J.A.G.; Marchioro, V.S.; Souza, V.Q.; Jost, E. Mean performance and stability in multi-environment trials I: Combining features of AMMI and BLUP techniques. Agron. J. 2019, 111, 2949–2960. 10.2134/agronj2019.03.0220.

Paris, B.; Vandorou, F.; Balafoutis, A.T.; Vaiopoulos, K.; Kyriakarakos, G.; Manolakos, D.; Papadakis, G. Energy use in greenhouses in the EU: A review recommending energy efficiency measures and renewable energy sources adoption. Appl. Sci. 2022, 12, 5150. 10.3390/app12105150.

Pezeshkpour, P.; Naseri, B.; Amiri, R.; Mirzaei, A.; Shobeiri, S.S.; Karami, I. Evaluation of mean performance and stability of lentil genotypes according to combination of additive main effects and multiplicative interaction, and best linear unbiased prediction methods. Legum. Sci., 2025, 7, e70021. 10.1002/leg3.70021.

Piepho, H.P. Best Linear Unbiased Prediction (BLUP) for regional yield trials: A comparison to additive main effects and multiplicative interaction (AMMI) analysis. Theor. Appl. Genet. 1994, 89, 647–654. 10.1007/BF00222462.

Piepho, H.P. Methods for comparing the yield stability of cropping systems—A review. J. Agron. Crop Sci. 1998, 180, 193–213. 10.1111/j.1439-037X.1998.tb00526.x.

Poore, J.; Nemecek, T. Reducing food’s environmental impacts through producers and consumers. Science 2018, 360, 987–992. 10.1126/science.aaq0216.

Preiti, G.; Calvi, A.; Badagliacca, G.; Lo Presti, E.; Monti, M.; Bacchi, M. Agronomic performances and seed yield components of lentil (*Lens culinaris* Medikus) germplasm in a semi-arid environment. Agronomy, 2024, 14, 303. 10.3390/agronomy14020303.

Rocchetti, L.; Gioia, T.; Logozzo, G.; Brezeanu, C.; Pereira, L.G.; De la Rosa, L.; Marzario, S.; Pieri, A.; Fernie, A.R.; Alseekh, S.; Susek, K.; Cook, D.R.; Varshney, R.K.; Agrawal, S.K., Hamwieh, A.; Bitocchi, E.; Papa, R. Towards the development, maintenance and standardized phenotypic characterization of single-seed-descent genetic resources for chickpea. Curr. Protoc. 2022, 2, e371. 10.1002/cpz1.371.

Ruisi, P.; Longo, M.; Martinelli, F.; Di Miceli, G.; Frenda, A.S.; Saia, S.; Carimi, F.; Giambalvo, D.; Amato, G. Morpho-agronomic and genetic diversity among twelve Sicilian agro-ecotypes of lentil (*Lens culinaris*). J. Anim. Plant Sci. 2015, 25, 716–728.

Sellami, M.H.; Pulvento, C.; Lavini, A. Selection of suitable genotypes of lentil (*Lens culinaris* Medik.) under rainfed conditions in south Italy using multi-trait stability index (MTSI). Agronomy 2021, 11, 1807. 10.3390/agronomy11091807.

Shobeiri, S.S.; Pezeshkpour, P.; Naseri, B. Evaluation of efficiency of weighted average of stability and mean performance estimated by linear mixed models for identifying high-yielding lentil genotypes adapted to rainfed regions. Legum. Sci., 2024a, 6, e226. 10.1002/leg3.226.

Shobeiri, S.S.; Pezeshkpour, P.; Naseri, B. Evaluation of seed yield stability of lentil genotypes by linear mixed effects models and multitrait stability index. Legum. Sci., 2024b, 6, e245. 10.1002/leg3.245

Shukla, P.R.; Skea, J.; Calvo Buendia, E.; Masson-Delmotte, V.; Pörtner, H.O.; Roberts, D.C.; Zhai, P.; Slade, R.; Connors, S.; Van Diemen, R.; Ferrat, M.; Haughey, E.; Luz, S.; Neogi, S.; Pathak, M.; Petzold, J.; Portugal Pereira, J.; Vyas, P.; Huntley, E.; Kissick, K.; Belkacemi, M.; Malley, J. Climate change and land: an IPCC special report on climate change, desertification, land degradation, sustainable land management, food security, and greenhouse gas fluxes in terrestrial ecosystems; 2019, Cambridge University Press, Cambridge, UK and New York, NY, USA, 896 pp. 10.1017/9781009157988.

Siddique, K.H.M.; Loss, S.P.; Pritchard, D.L.; Regan, K.L.; Tennant, D.; Jettner, R.L.; Wilkinson, D. Adaptation of lentil (*Lens culinaris* Medik.) to Mediterranean-type environments: Effect of time of sowing on growth, yield, and water use. Austr. J. Agric. Res. 1998, 49, 613–626. 10.1071/A97128.

Silva-Perez, V.; Shunmugam, A. S.; Rao, S.; Cossani, C. M.; Tefera, A. T.; Fitzgerald, G. J.; Armstrong, R.; Rosewarne, G. M. Breeding has selected for architectural and photosynthetic traits in lentils. Front. Plant Sci., 2022, 13, 925987. 10.3389/fpls.2022.925987.

Sonnante, G.; Hammer, K.; Pignone, D. From the cradle of agriculture a handful of lentils: History of domestication. Rend. Lincei. Sci. Fis. Nat. 2009, 20, 21–37. 10.1007/s12210-009-0002-7.

Summerfield, R.J.; Roberts, E.H.; Erskine, W.; Ellis, R.H. Effects of temperature and photoperiod on flowering in lentils (*Lens culinaris* Medic.). Ann. Bot., 1985, 56, 659–671. 10.1093/oxfordjournals.aob.a087055.

Teichmann, T.; Muhr, M. Shaping plant architecture. Front. Plant Sci. 2015, 6, 233. 10.3389/fpls.2015.00233.

Van Eeuwijk, F.A. Linear and bilinear models for the analysis of multi-environment trials: I. An inventory of models. Euphytica 1995, 84, 1–7. 10.1007/BF01677551.

Watson, C.A.; Reckling, M.; Preissel, S.; Bachinger, J.; Bergkvist, G.; Kuhlman, T.; Lindström, K.; Nemecek, T.; Topp, C.F.E.; Vanhatalo, A. Grain legume production and use in European agricultural systems. In Advances in Agronomy; Elsevier: Amsterdam, The Netherlands, 2017; Volume 144, pp. 235–303.

Wright, D.M.; Neupane, S.; Heidecker, T.; Haile, T.A.; Chan, C.; Coyne, C.J.; McGee, R.J.; Udupa, S.; Henkrar, F.; Barilli, E.; Rubiales, D.; Gioia, T.; Logozzo, G.; Marzario, S.; Mehra, R.; Sarker, A.; Dhakal, R.; Babul Anwar, B.; Sarkar, D.; Vandenberg, A.; Bett, K.E. Understanding photothermal interactions will help expand production range and increase genetic diversity of lentil (*Lens culinaris* Medik.). Plants People Planet 2021, 3, 171–181. 10.1002/ppp3.10158.

Yan, W. GGEbiplot-A Windows application for graphical analysis of multi-environment trial data and other types of two-way data. Agron. J. 2001, 93, 1111–1118. 10.2134/agronj2001.9351111x.

Zaccardelli, M.; Lupo, F.; Piergiovanni, A.R.; Laghetti, G.; Sonnante, G.; Daminati, M.G.; Sparvoli, F.; Lioi, L. Characterization of Italian lentil (*Lens culinaris* Medik.) germplasm by agronomic traits, biochemical and molecular markers. Genet. Resour. Crop Evol. 2012, 59, 727–738. 10.1007/s10722-011-9714-5.

Zeroual, A.; Mitache, M.; Baidani, A.; Abderemane, B.A.; Benbrahim, N.; Ouhemi, H.; Çakır, E.; Hoyos-Villegas, V.; Gadaleta, A.; Mazzucotelli, E.; Özkan, H.; Idrissi, O. Assessment of the phenotypic diversity and agronomic performance of a Mediterranean lentil collection under rainfed conditions: towards efficient use in breeding programs for adaptation to Mediterranean-type environment. Genet. Resour. Crop Evol., 2024, 10.1007/s10722-024-02115-y.

Zohary, D. The wild progenitor and the place of origin of the cultivated lentil: *Lens culinaris*. Econ. Bot. 1972, 26, 326–332. 10.1007/BF02860702.

